# *Prdm16* mutation determines sex-specific cardiac metabolism and identifies two novel cardiac metabolic regulators

**DOI:** 10.1101/2023.01.10.523243

**Authors:** Jirko Kühnisch, Simon Theisen, Josephine Dartsch, Raphaela Fritsche-Guenther, Marieluise Kirchner, Benedikt Obermayer, Anna Bauer, Anne-Karin Kahlert, Michael Rothe, Dieter Beule, Arnd Heuser, Philipp Mertins, Jennifer A. Kirwan, Nikolaus Berndt, Calum A. MacRae, Norbert Hubner, Sabine Klaassen

**Affiliations:** Experimental and Clinical Research Center, a cooperation between the Max Delbrück Center for Molecular Medicine in the Helmholtz Association and Charité - Universitätsmedizin Berlin, Germany; Max Delbrück Center for Molecular Medicine in the Helmholtz Association (MDC), Berlin, Germany; DZHK (German Centre for Cardiovascular Research), partner site Berlin, Berlin, Germany; Berlin Institute of Health (BIH) at Charité - Universitätsmedizin Berlin, BIH Metabolomics Platform, Berlin, Germany; Max-Delbrück-Center for Molecular Medicine in the Helmholtz Association (MDC), Proteomics Platform, Berlin, Germany; Berlin Institute of Health (BIH) at Charité - Universitätsmedizin Berlin, Berlin, Germany; Berlin Institute of Health at Charité - Universitätsmedizin Berlin, Core Unit Bioinformatics, Berlin, Germany; Department of Congenital Heart Disease and Pediatric Cardiology, University Hospital of Schleswig-Holstein, Kiel, Germany; German Center for Cardiovascular Research (DZHK), Kiel, Germany; Institute of Immunology and Genetics, Kaiserslautern, Germany; Lipidomix GmbH, Berlin, Germany; Institute of Computer-Assisted Cardiovascular Medicine, Charité - Universitätsmedizin Berlin, corporate member of Freie Universität Berlin and Humboldt-Universität zu Berlin, Berlin, Germany; Harvard Medical School and Cardiovascular Division, Department of Medicine, Brigham and Women’s Hospital, Boston, USA; Department of Pediatric Cardiology, Charité - Universitätsmedizin Berlin, corporate member of Freie Universität Berlin and Humboldt-Universität zu Berlin, Berlin, Germany

## Abstract

**Background:** Mutation of the *PRDM16* gene has been associated with human cardiomyopathy. The PRDM16 protein is a transcriptional regulator affecting cardiac development via Tbx5 and Hand1 regulating myocardial structure. Biallelic *Prdm16* inactivation induces severe cardiac dysfunction with postnatal lethality and hypertrophy in mice. Early pathological events upon *Prdm16* inactivation have not been explored.

**Methods:** This study performed in depth pathophysiological and molecular analysis of male and female *Prdm16^csp1/wt^* mice carrying systemic, monoallelic *Prdm16* gene inactivation. We systematically assessed early molecular changes with transcriptomics, proteomics, and metabolomics. Kinetic modelling of the cardiac metabolism was undertaken *in silico* with CARDIOKIN.

**Results:** *Prdm16^csp1/wt^* mice are viable up to 8 months, develop hypoplastic hearts, and diminished systolic performance that is more pronounced in female mice. *Prdm16^csp1/wt^* hearts demonstrate moderate alterations of specific transcripts and protein levels with consistent upregulation of pyridine nucleotide-disulphide oxidoreductase domain 2 (Pyroxd2) and the transcriptional regulator pre B-cell leukemia transcription factor interacting protein 1 (Pbxip1). The strongest concordant transcriptional upregulation was detected for *Prdm16* itself probably by an autoregulatory mechanism. *Prdm16^csp1/wt^* cardiac tissue showed reduction of metabolites associated with amino acid as well as glycerol metabolism, glycolysis, and tricarboxylic acid cycle. Global lipid metabolism was also affected with accumulation of triacylglycerides detected in male *Prdm16^csp1/wt^* hearts. In addition, *Prdm16^csp1/wt^* cardiac tissue revealed diminished glutathione (GSH) and increased inosine monophosphate (IMP) levels indicating oxidative stress and a dysregulated energetics, respectively. Metabolic modelling *in silico* suggested lowered fatty acid utilization in male and reduced glucose utilization in female *Prdm16^csp1/wt^* cardiac tissue.

**Conclusions:** Monoallelic *Prdm16* mutation restricts cardiac performance in *Prdm16^csp1/wt^* mice. Metabolic alterations precede transcriptional dysregulation in *Prdm16^csp1/wt^* cardiac tissue. Female *Prdm16^csp1/wt^* mice develop a more pronounced phenotype indicating a sexual dimorphism at this early pathological window. This study suggests that metabolic dysregulation is an early event in *PRDM16* associated cardiac pathology.

**Novelty and Significance:** *What Is Known?:*

- Mutation of the *PRDM16* gene has been associated with human cardiomyopathy.
- Biallelic inactivation of *Prdm16* in mice induces severe cardiac dysfunction with early postnatal lethality.
- Prdm16 cooperates with transcription factors such as Tbx5 and Hand1 to activate transcriptional programs that define the development of the compacted myocardium.

*What New Information Does This Article Contribute?:*

- Systemic, monoallelic inactivation in *Prdm16^csp1/wt^* mice induces cardiac dysfunction with normal survival.
- Metabolic alterations are the leading pathophysiological consequences and induce cardiac hypoplasia. On the molecular level this is associated with upregulation of metabolic regulators Pyroxd2 and Pbxip1.
- Metabolic response after *Prdm16* inactivation occurs in a sex specific manner.

## Background

Primary, genetically determined cardiomyopathies comprise a group of heterogenous cardiac diseases that eventually result in heart failure or arrhythmia. Approximately 100 genes have been linked to cardiomyopathy most frequently affecting the sarcomere, Z-disc, mitochondria, or ion channel regulation.^1,2^ In addition to these disease circuits transcription and splicing may be disturbed in cardiomyopathy. The PR/SET domain 16 (PRDM16) protein is one such transcriptional regulator.^3^ Mutation of the *PRDM16* gene causes dilated (DCM) and left ventricular noncompaction cardiomyopathy (LVNC) in patients.^4–7^ Recently, rare variant association analysis linked LVNC to PRDM16 protein truncating variants.^8^ Altogether, this suggests that *PRDM16* is a genetic factor critical for cardiac function either causing monogenic cardiomyopathy or other myocardial phenotypes.

Germline homozygous inactivation of *Prdm16* in *Prdm16^csp1/csp1^* mice results in a complex phenotype involving several organs, cardiac hypoplasia, and early postnatal lethality.^9^ More recently, the role of *Prdm16* in myocardial development and LVNC was assessed after homozygous, cardiac-specific *Prdm16* inactivation in *Xmlc2Cre;Prdm16^flox/flox^* mice.^10^ Homozygous *Xmlc2Cre;Prdm16^flox/flox^* mice develop cardiac dysfunction and die prematurely before postnatal day 7. During cardiac development, Prdm16 cooperates with the transcription factors T-box 5 (Tbx5) and heart and neural crest derivatives expressed 1 (Hand1) to promote gene programs required for myocardial growth and compaction.^10^ Prdm16 also suppresses neural gene expression.^10^ Consequently, Prdm16 inactivation in *Xmlc2Cre;Prdm16^flox/flox^* mice leads to biventricular hypertrabeculation and left ventricular dilatation.^10^ Overall, this work established that *Prdm16* is critical for embryonic cardiac development and postnatal function, orchestrating the transcriptional circuits determining myocardial maturation.

Originally, PRDM16 was established as a determinant for differentiation, homeostasis, and function of brown/beige adipocytes.^11^ PRDM16 induces gene programs for the development of brown adipocytes, represses muscle/white adipocyte specific genes, induces adaptive thermogenesis, and increases energy expenditure.^12–15^ On a molecular level these effects are facilitated by physical interaction and coordination of key transcription factors such as peroxisome proliferator activated receptor alpha, gamma (PPARA, PPARG), mediator complex subunit 1 (MED1), or CCAAT enhancer binding protein delta (CEBPD).^12,16,17^ Of note, a significant number of these PRDM16 associated proteins serve as critical regulators of fatty acid (FA) metabolism, glucose utilization, and/or cellular respiration.

This study tests the hypothesis that PRDM16 orchestrates cardiac metabolism and investigates its role beyond its known transcriptional functions in cardiac development and homeostasis. We characterize the heart of heterozygous *Prdm16^csp1/wt^* mice in depth to assess the impact of monoallelic germline *Prdm16* inactivation, as is present in patients with *PRDM16* associated cardiomyopathy. We explore early molecular events upon *Prdm16* inactivation and establish a preclinical animal model. *Prdm16^csp1/wt^* mice show diminished cardiac performance, normal survival, and altered body composition. Cardiac dysfunction is explained on the molecular level by altered metabolism, redox balance, and FA/glucose utilization. Overall, this study establishes heterozygous *Prdm16^csp1/wt^* mice as a model for early molecular pathomechanistic events in the development of the *PRDM16* associated cardiomyopathy.

## Methods

The high-throughput sequencing data and proteome data have been made publicly available at the Gene Expression Omnibus (GEO accession No. …) and the Proteomics Identification Database (PRIDE accession No. …). Other data and study materials are available from the corresponding authors on reasonable request. Primer and antibodies used in this study are available (Table I and II in the Data Supplement). A detailed material and methods section is provided in the Supplemental Material. The *Prdm16^csp1/wt^* mice (FVB.C-*Prdm16^csp1^*/J) were received from Jackson Laboratories, USA (JAX stock #013100). The FVB.C-*Prdm16^csp1^*/J strain was originally established by N-ethyl-N-nitrosourea (ENU) mutagenesis inducing a missense C>A mutation at the intronic acceptor splice site of exon 7.^9^ Maintenance, physiological analysis and organ collection of *Prdm16^csp1/wt^* mice was approved by the Landesamt für Gesundheit und Soziales Berlin (LAGeSo), Germany (G0070/17).

## Results

### Germline, heterozygous inactivation of Prdm16 induces mild cardiac dysfunction

In human tissue RNA preparations *PRDM16* transcripts are most abundant in lung, followed by aorta, adipose tissues, and heart (Figure 1A). Consistently, murine tissue RNA extracts show approx. 10-fold higher *Prdm16* expression in lung compared to heart (Figure 1B). Within different heart regions *Prdm16* shows abundant expression in the right ventricle (RV), left ventricle (LV), and septum but not in the atria (Figure 1C). Potassium voltage-gated channel member 4 (*Kcna4*) and myosin light chain 2 (*Myl2*) confirmed atrial and ventricular origin, respectively. To assess clinically relevant physiological and molecular impact of *Prdm16* in the heart, rather than the effects of biallelic *Prdm16* gene inactivation, we analyzed heterozygous FVB.C-*Prdm16^csp1^/J* mice (*Prdm16^csp1/wt^*).^9^ PCR genotyping and Sanger sequencing confirmed presence of the c.888-3C>A (ENSMUSG00000039410) variant on DNA level (Figure 1D, Figure I_A-B in the Data Supplement). To further validate the impact of the *Prdm16* acceptor splice site variant c.888-3C>A (ENSMUST00000030902.12) at the transcriptional level, we performed PCR and targeted high throughput sequencing of total RNA isolated from different *Prdm16^wt/wt^* and *Prdm16^csp1/wt^* tissues. These analyses suggest that the *Prdm16* acceptor splice site variant c.888-3C>A affects mRNA splicing, produces several splice products, and the most abundant splice products truncate Prdm16 proteins after approx. 340 amino acids (Figure 1E-F, for detailed Results see Data Supplement).

**Figure 1.**
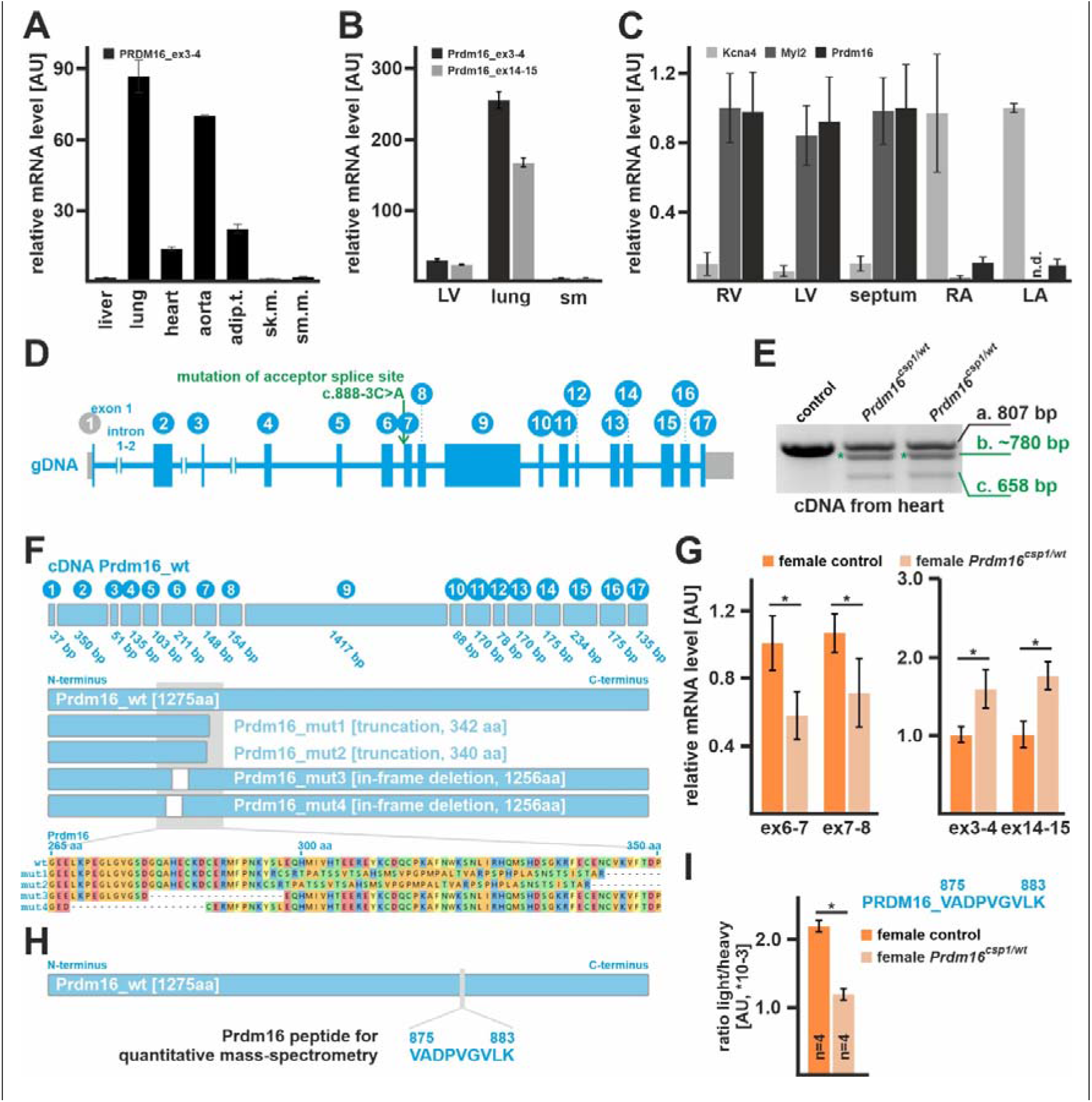
Expression of *Prdm16* in the heart of *Prdm16^csp1/wt^* mice. **(A)** Expression analysis reveals high *PRDM16* transcript levels in human lung, aorta, adipose tissue (adip.t.), and heart. Liver, skeletal muscle (sk.m.), and smooth muscle (sm.m.) show low *PRDM16* expression. **(B)** *Prdm16* is highly expressed in murine lung tissue, shows robust levels in the left ventricle (LV) but low expression in skeletal muscle (sm). **(C)** *Prdm16* is robustly expressed in LV, right ventricle (RV), and the septum. The left (LA) and right atrium (RA) does not show considerable *Prdm16* expression. *Kcna4* and *Myl2* transcripts demonstrate atrial and ventricular tissue origin. **(D)** The murine *Prdm16* gene comprises 17 exons and *Prdm16^csp1/wt^* mice carry the point mutation c.888-3C>A affecting the acceptor splice site of intron_6-7. **(E)** PCR genotyping with heart cDNA from heterozygous *Prdm16^csp1/wt^* mice generates 3 products: a) wild-type fragment (807 bp), b) unknown mutant product (~780 bp, *), and c) mutant product without exon 7 (658 bp). **(F)** The wildtype Prdm16 protein is 1275 amino acids (aa) long. Targeted high-throughput sequencing of the different PCR products identifies four main Prdm16 mutant proteins in *Prdm16^csp1/wt^* mice including truncation (mut1, mut2) and in-frame deletion (mut3, mut4) variants (Figure I and II in the Data Supplement, for detailed Results see Data Supplement). **(G)** Quantitative detection of *Prdm16* transcript levels with qPCR detectors targeting the exon 7 deletion region shows approx. 40-50% reduced expression. In contrast, PCR detectors targeting exon3-4 and exon14-15 identifies increased *Prdm16* levels. Analysis was performed on female *Prdm16^csp1/wt^* heart tissue. **(H)** Detection of Prdm16 protein was achieved by PRM (parallel-reaction monitoring) using the Prdm16 derived peptide V875- K883. **(I)** Quantitation of Prdm16 peptide in control and *Prdm16^csp1/wt^* lung tissue. The ratio of endogenous (light) and standard spike-in (heavy) peptide is shown. Statistical analysis was performed with unpaired t-test (p<0.05).

Next, we tested *Prdm16* mutant transcript expression in the heart. *Prdm16* mutant transcripts are ~40% diminished compared to controls using primer sets targeting the mutated exon6-8 region (Figure 1G, Figure I_F in the Data Supplement). In contrast, primer sets measuring the overall *Prdm16* transcript level (exon3-4, exon14-15) detect increased expression in *Prdm16^csp1/wt^* hearts. This suggests that *Prdm16* inactivation due to the c.888-3C>A variant upregulates overall *Prdm16* mRNA expression. The Prdm16 protein level was assessed with a mass spectrometry-based targeted proteomics approach, measuring and quantifying the abundance of the peptide Prdm16_V875-K883 (Figure 1H). As expected, a ~50% reduction of the Prdm16_V875-K883 level was observed in *Prdm16^csp1/wt^* lung compared to controls (Figure 1I). In cardiac tissue the Prdm16_V875-K883 detection limit was not reached.

The survival of *Prdm16^csp1/wt^* mice is normal until 8 months (Figure 2A). Male and female *Prdm16^csp1/wt^* mice exhibit body weight reduction by ~11% and ~20%, respectively (Figure 2B, Table III in the Data Supplement). In female *Prdm16^csp1/wt^* mice diminished body weight was associated with reduced fat content (~14%) and increased relative muscle tissue content (~8%). The absolute and relative heart weight is diminished in both male and female *Prdm16^csp1/wt^* mice (Figure 2B). Transthoracic echocardiography was assed to characterize the cardiac physiology of *Prdm16^csp1/wt^* mice. Echocardiography illustrates mild cardiac hypoplasia and reduced systolic functional parameters such as stroke volume, cardiac output, and ejection fraction (EF) (Figure 2C-D, Table IV-VI in the Data Supplement). Other cardiac parameters such as heart rate, systolic/diastolic blood pressure, or electrophysiology are unaffected in *Prdm16^csp1/wt^* mice. Cardiac tissue organization assessed by histology and hematoxylin/eosin staining appeared normal in *Prdm16^csp1/wt^* hearts (Figure 2E). Fibrosis was not detected, neither by Picro-Sirius red staining, nor by immunostaining of collagen 1 (Col1) and alpha smooth muscle actin (αSma) (Figure 2F, Figure III in the Data Supplement). Further analysis of *Prdm16^csp1/wt^* cardiac tissue with electron microscopy, morphometry, quantitative PCR, and immunostaining identified no abnormalities of the sarcomere (Figure IV in the Data Supplement, for detailed Results see Data Supplement). The relative mitochondrial area in heart tissue was unaffected. Cardiomyocyte area was assessed histomorphometrically in paraffin embedded heart tissue sections of comparable cross-sectional level after wheat germ agglutinin (WGA) staining. Cardiomyocytes of female *Prdm16^csp1/wt^* mice demonstrate significant reduction of cross-sectional area explaining reduced myocardial mass (Figure 2G). Normal viability, myocardial hypoplasia, and diminished cardiac performance due to monoallelic *Prdm16* inactivation suggests *Prdm16^csp1/wt^* mice as a suitable model to explore early pathophysiological changes in *PRDM16* associated cardiomyopathy.

**Figure 2.**
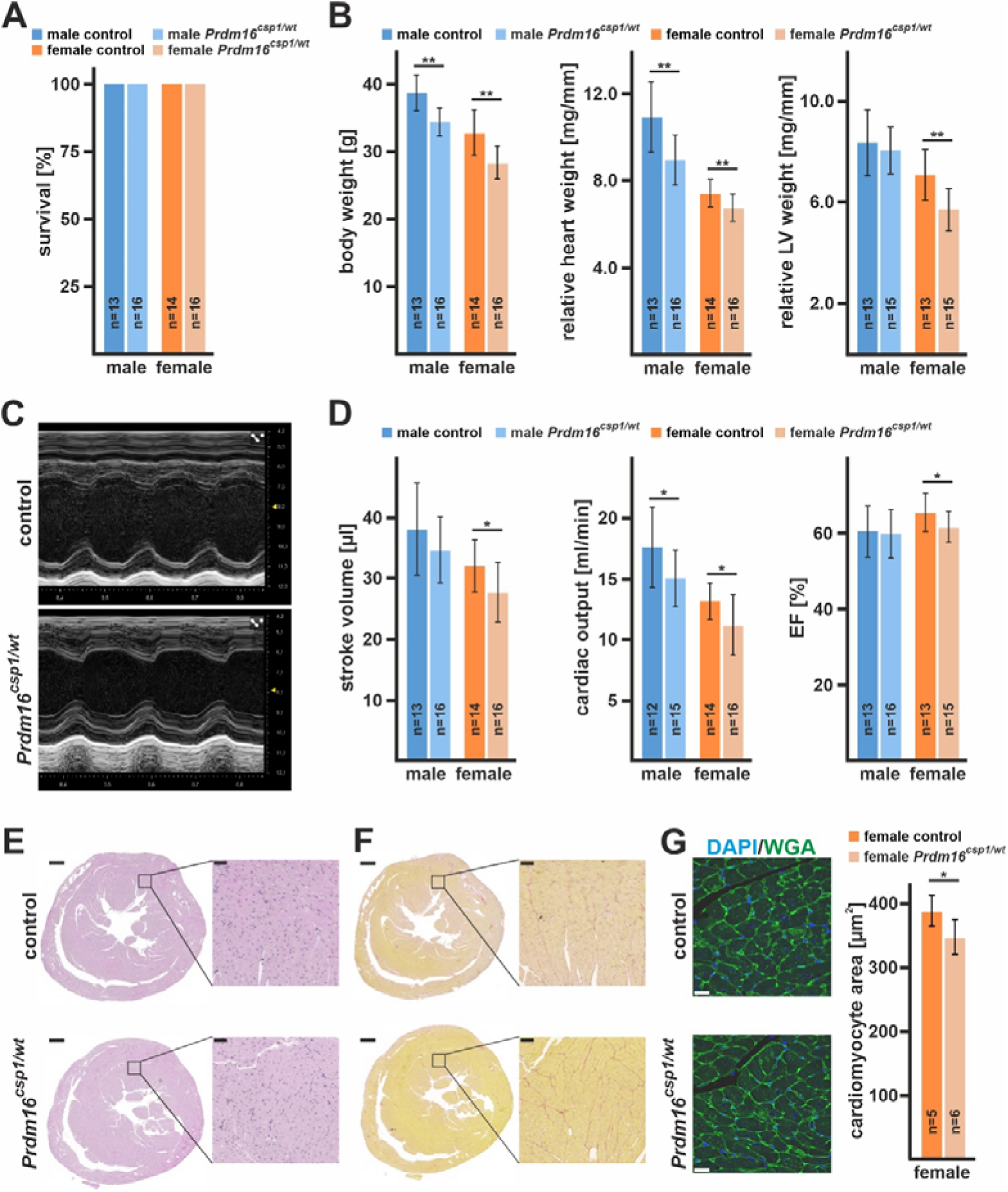
Diminished cardiac performance after *Prdm16* inactivation. **(A)** Survival of male and female *Prdm16^csp1/wt^* mice is normal until the age of 8 month. **(B)** 8-month-old *Prdm16^csp1/wt^* mice of both sexes have a reduced absolute body weight. Normalization against tibia length demonstrated a diminished relative heart and LV weight in *Prdm16^csp1/wt^* mutants. **(C)** M-mode images illustrate the heart phenotype in *Prdm16^csp1/wt^* mice as assessed with echocardiography. **(D)** At the age of 8 months *Prdm16^csp1/wt^* mice of both sexes have diminished cardiac function as illustrated by stroke volume, cardiac output, and ejection fraction (EF). Full echocardiography data are available Tables IV-VI in the Data Supplement. **(E)** Histological analysis with H&E staining reveals normal tissue organization in female *Prdm16^csp1/wt^* hearts. Scale bars in E and F are 500 μm (full heart) and 50 μm (inset). **(F)** Histological staining for fibrosis with Picro-Sirius red is negative in female *Prdm16^csp1/wt^* heart tissue. **(G)** The cell area of female heart muscle (cross sections) stained with wheat-germ agglutinin (WGA, green) is significantly reduced. Nuclei were stained with DAPI (blue). Scale bar is 20 μm. Statistical analysis was performed with unpaired t-test (p<0.05).

### Moderate transcriptional dysregulation in Prdm16^csp1/wt^ hearts

Prdm16 is a transcriptional regulator so we performed transcriptional profiling using RNAseq of heart tissue comparing male and female *Prdm16^csp1/wt^* mice with corresponding controls. Comparing female to male control heart tissue, we found 51 up-regulated and 92 down-regulated genes, applying an absolute log2 fold change (LFC) threshold >0.5 (Figure 3A). Male *Prdm16^csp1/wt^* mice showed 12 up-regulated and 16 down-regulated genes compared to male controls. Female *Prdm16^csp1/wt^* mice revealed 55 up-regulated and 35 down-regulated genes, respectively. Transcriptional profiling identified most significant up-regulation of *Prdm16* in males and females suggesting an autoregulatory control of expression (Figure 3B-C). Other transcripts of the PRDM gene family were not regulated (Figure V_A-B in the Data Supplement).

**Figure 3.**
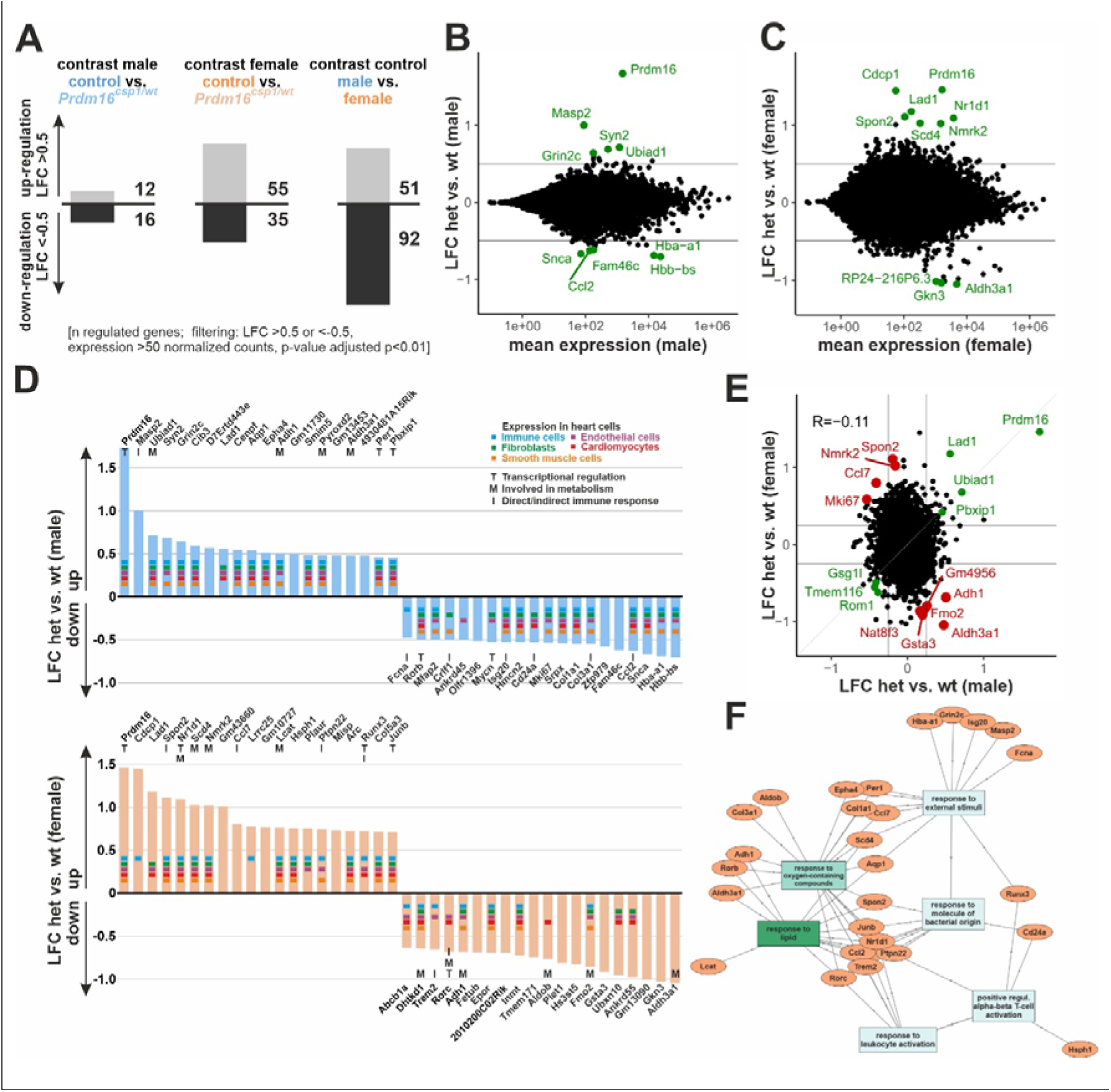
Transcriptome analysis of *Prdm16^csp1/wt^* heart tissue. **(A)** Transcriptome analysis with RNAseq of *Prdm16^csp1/wt^* LV shows in male 28 and in female 90 differentially regulated genes compared to controls. Analysis of male vs. female control hearts identifies 143 differentially regulated transcripts. **(B-C)** Bioinformatic filtering ranked the normalized log2 cpk according the absolute log2 fold change (LFC) after elimination of regulated targets with an adjusted p-value (padj) <10^-2^. Scatterplots show the LFC against mean expression for the contrast *Prdm16^csp1/wt^* (het) vs controls (wt) in male (B) or female (C) heart. The top 10 differentially regulated genes (adj. p < 0.01) are highlighted in green and labeled. **(D)** Summary of top 20 up- and down-regulated genes in male (upper panel) and female (lower panel) hearts. Expression of selected genes in cardiac cell types is annotated according to the color code. Functional association with transcription (T), metabolism (M), and immune response (I) is shown for relevant genes. **(E)** Scatterplot shows the LFC detected in A and B against each other. Genes with concordant changes (abs LFC > 0.25) are highlighted in green, genes with discordant changes (top 10; adj. p < 0.01) are highlighted in red. The overall correlation is R=-0.11 (n=21276). **(F)** Gene ontology (GO) network was constructed with GOnet^19^ by using the male and female TOP20 up- and down-regulated genes in *Prdm16^csp1/wt^* hearts. GO terms *response to lipids* and *response to oxygen-containing compounds* show strongest association.

Next, the top 20 up- and down-regulated genes for males and females were evaluated. In male *Prdm16^csp1/wt^* mice we observed upregulation for MBL associated serine protease 2 (*Masp2*), synapsin II (*Syn2*) a coat protein of clathrin-coated vesicles, and UbiA prenyltransferase domain containing 1 (*Ubiad1*) controlling coenzyme Q10 synthesis (Figure 3D). We detected decreased levels of transcripts for synuclein alpha (*Snca*), hemoglobin alpha adult chain 1 (*Hba-a1*), and hemoglobin subunit beta (*Hbb-ba*), which both supply/control intracellular oxygen. Female *Prdm16^csp1/wt^* cardiac tissue showed up-regulation of the extra cellular matrix CUB domain containing protein 1 (*Cdcp1*), ladinin (*Lad1*) involved in basement membrane organization, the cell adhesion protein spondin 2 (*Spon2*), nuclear receptor subfamily 1, group D, member 1 (*Nr1d1*) a transcriptional repressor coordinating metabolic pathways, and nicotinamide riboside kinase 2 (*Nmrk2*) regulating laminin matrix deposition. Aldehyde dehydrogenase family 3, subfamily A1 (*Aldh3a1*) and gastrokine 3 (*Gkn3*) showed strongest down-regulation in female *Prdm16^csp1/wt^* cardiac tissue. The top 20 genes were tested for their cellular expression in heart tissue (www.proteinatlas.org). Most genes showed a broad expression in cardiomyocytes, endothelial cells, fibroblasts, immune cells, and smooth muscle cells. Thus, no enrichment of cell lineage specific genes was found. The majority of dysregulated genes in male and female *Prdm16^csp1/wt^* cardiac tissue are associated with transcription (T), metabolism (M), and indirect/direct immune response (I).

Concordant and discordant changes in males vs. females were assessed with LFC significance threshold of >0.25 and <-0.25, respectively. Strongest concordant up-regulated genes in *Prdm16^csp1/wt^* hearts were *Prdm16, Lad1, Ubiad1*, and the pre B-cell leukemia transcription factor interacting protein 1 (*Pbxip1*) (Figure 3E). Systematic dysregulation of critical cardiac genes was tested using the harmonizome gene set *congenital heart disease*. Evaluation did not identify consistent, significant dysregulation of cardiac specific transcripts (Figure V_C in the Data Supplement). Differentially expressed genes from *Prdm16^csp1/wt^* hearts (TOP5 up- and down-regulated genes) and known PRDM16 interacting genes (physical interaction, transcriptional targets, transcriptional complex component) were tested for dysregulation in a human DCM patients dataset.^18^ From more than 60 known PRDM16 targets *PPARA, MED1*, and *CEBPD* were dysregulated in the human DCM screen (Table VII in the Data Supplement). Moreover, the chemotactic factor 2 (*Ccl2*) was dysregulated in DCM patients and in male *Prdm16^csp1/wt^* mice. Sex specific gene expression validated correct technical procedures of RNAseq *Prdm16^csp1/wt^* mice analysis (Figure V_D in the Data Supplement). Next, we generated a gene ontology (GO) network using the male and female TOP20 up- and down-regulated genes in *Prdm16^csp1/wt^* hearts (GOnet).^19^ Strongest functional association of dysregulated genes in *Prdm16^csp1/wt^* hearts was found for the GO terms *response to lipids* and *response to oxygen-containing compounds* (Figure 3F, Figure V_E in the Data Supplement). Thus, *Prdm16^csp1/wt^* hearts show moderate, sex specific transcriptional dysregulation with the strongest regulated transcripts being *Prdm16, Lad1, Ubiad1*, and *Pbxip1*. GO and individual transcript evaluation suggest an impact on metabolic processes after monoallelic *Prdm16* inactivation.

### Proteome analysis of Prdm16^csp1/wt^ hearts reveals upregulation of Pbxip1 and Pyroxd2

In order to verify findings from the transcriptional analysis and to assess cardiac tissue protein expression, we performed a global proteomic analysis of cardiac tissue. Consistent with RNAseq, male *Prdm16^csp1/wt^* cardiac tissue showed less variation from control than female samples. Of the 3,847 proteins identified in total, 3,314 were used for quantitation. Sample identity was confirmed by expression level of the male-specific DEAD box helicase 3, Y-linked (Ddx3y) and eukaryotic translation initiation factor 2, subunit 3, structural gene Y-linked (Eif2s3y). The Prdm16 protein was not detected, which can be explained by very low abundance below the limit of detection. Most significant differences in protein expression were observed between male and female cardiac tissue (165 proteins with FDR <5%, data not shown). Pairwise evaluation of control and *Prdm16^csp1/wt^* proteome data from male cardiac tissue using a p-value cutoff 0.01 showed upregulation of fermitin family member 2 (Fermt2), coatomer protein complex, subunit zeta 2 (Copz2), and pyridine nucleotide-disulphide oxidoreductase domain 2 (Pyroxd2) (Figure 4A). Substantial downregulation was observed for c-src tyrosine kinase (Csk), translocase of inner mitochondrial membrane 10B (Timm10b), and collagen type XVIII, alpha 1 (Col18a1) proteins. Pairwise evaluation of proteome data from female cardiac tissue showed strongest upregulation of Pyroxd2, pyruvate dehydrogenase kinase, isoenzyme 4 (Pdk4), and pre B cell leukemia transcription factor interacting protein 1 (Pbxip1) in *Prdm16^csp1/wt^* mice (Figure 4B). Strongest downregulation was detected for cystic fibrosis transmembrane conductance regulator (Cftr), WNK lysine deficient protein kinase 1 (Wnk1), and huntingtin interacting protein 1 (Hip1).

**Figure 4.**
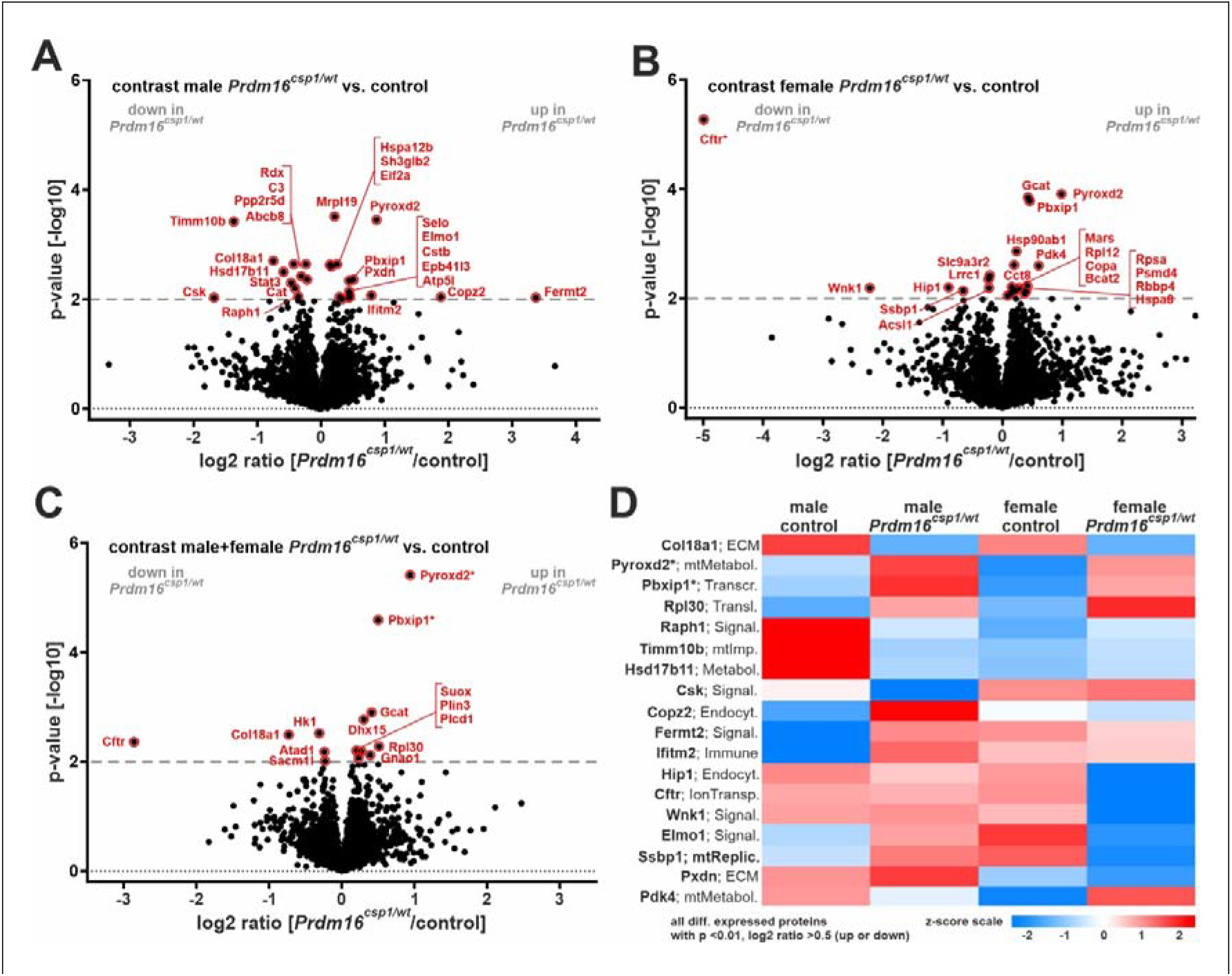
Proteome comparison of *Prdm16^csp1/wt^* cardiac tissue. Volcano plots for *Prdm16^csp1/wt^*/control pairwise comparisons with log2 ratio (x-axis) and p-value (y-axis) of male hearts **(A)**, female hearts **(B)**, and combined analysis of male+female hearts **(C)**. Red coloring indicates significantly altered proteins (threshold dashed grey line, p-value <0.01) and * highlights highly significantly different proteins with p-value <0.0001. **(D)** Heat map of all regulated proteins (p-value <0.01, abs log2 ratio >0.5) using z-scored values of group median intensity. Pre B-cell leukemia transcription factor interacting protein 1 (Pbxip1) and pyridine nucleotide-disulphide oxidoreductase domain 2 (Pyroxd2) are consistently increased in male and female *Prdm16^csp1/wt^* mice with p-value <0.0001.

To further assess *Prdm16* genotype driven differences, we performed also pairwise evaluation of combined male and female groups (p-value cutoff 0.01). In total we identified 53 up- or downregulated proteins (Figure 4C, Figure VI in the Data Supplement). Pyroxd2, Pbxip1, and ribosomal protein L30 (Rpl30) appeared as the strongest concordantly upregulated proteins in *Prdm16^csp1/wt^* cardiac tissue. Furthermore, the strongest concordant downregulation was found for Cftr, Col18a1, and hexokinase 1 (Hk1) in *Prdm16^csp1/wt^* cardiac tissue. By adding an additional filter (abs log2 ratio >0.5), the top regulated proteins (n=18) were selected (Figure 4D). Among these, Pyroxd2 and Pbxip1 represent the two most significantly regulated candidates in *Prdm16^csp1/wt^* cardiac tissue (p-value<0.0001). Pyroxd2 is a critical regulator of hepatic mitochondrial function, interacts with mitochondrial complex IV, and appears consistently upregulated in *Prdm16^csp1/wt^* hearts on transcript as well as protein level.^20^ Pbxip1 interacts with transport as well as regulatory proteins and indirectly affects transcription.^21^ Proteome data did not show differential expression of mitochondrial transport proteins and respiratory chain complexes (Figure VII in the Data Supplement). Proteome data for sarcomere components, glycolysis, and amino acid metabolism did not show differential expression (Figure VIII in the Data Supplement). Altogether protein expression analysis identifies specific dysregulation of the mitochondrial protein Pyroxd2 and the transcriptional regulator Pbxip1.

### Altered metabolism in Prdm16^csp1/wt^ cardiac tissue

As expression analysis pointed towards a metabolic impact of Prdm16 in cardiac tissue, we also analyzed central carbon metabolites in *Prdm16^csp1/wt^* cardiac tissue with gas chromatography mass spectrometry (GC-MS). Measured values are presented as log2 from ratio of mean of the normalized peak areas *Prdm16^csp1/wt^*/controls (values >0.2, blue and values <-0.2, red). Overall, we detected several diminished metabolites (log2 ratios *Prdm16^csp1/wt^*/controls) for all assessed metabolic processes (Figure 5A). In *Prdm16^csp1/wt^* cardiac tissue, amino acid, glycerol, pentose phosphate pathway (PPP), glycolysis, tricarboxylic acid cycle (TCA), and nucleobase metabolism were all reduced. Individual metabolite analysis of female *Prdm16^csp1/wt^* cardiac tissue revealed statistically significant reductions for glycerol-3-phosphate, phosphoenolpyruvic acid, succinic acid, and 3-hydroxy butanoic acid. Combined female and male cardiac tissue analysis identified significant reduction of phosphoenolpyruvic acid, pyruvic acid, and ribose-5-phosphate in *Prdm16^csp1/wt^* hearts. As interpretation of individual metabolites is difficult, we assessed the designated pathway profiles of central carbon metabolites with univariate analysis. Univariate analysis counts each increased/diminished metabolite as ordinary number. Combined pathways analysis using female and male values identified significant reduction of amino acid, glycolysis, glycerol, and TCA metabolism in *Prdm16^csp1/wt^* hearts (Figure 5B).

**Figure 5.**
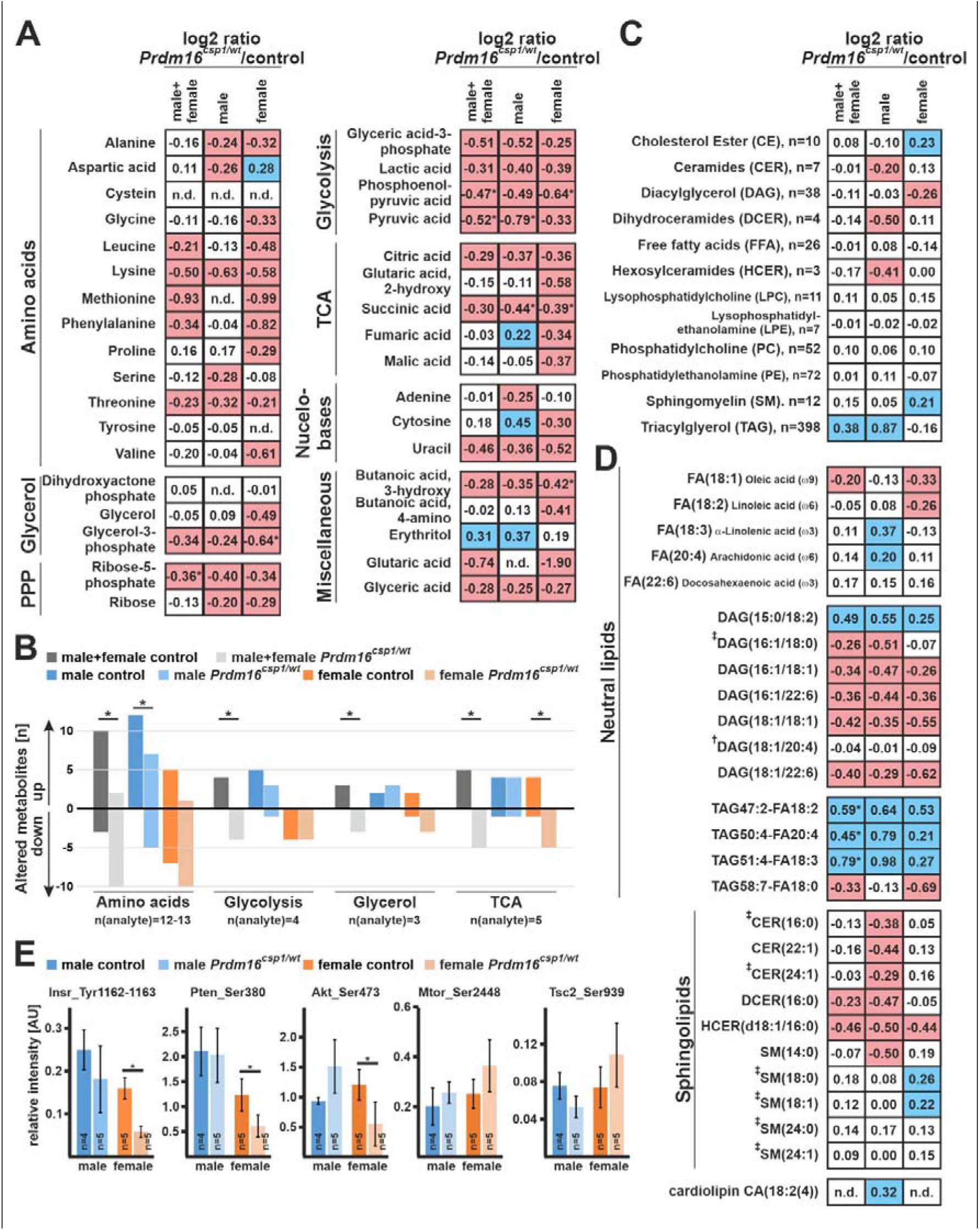
Altered metabolism in *Prdm16^csp1/wt^* cardiac tissue. **(A)** Normalized central carbon metabolite counts are presented as log2 value of the mean of normalized peak area ratio *Prdm16^csp1/wt^*/controls of males, females, and combination of both sexes. In female *Prdm16^csp1/wt^* LV tissue, broad suppression of several metabolic pathways was observed. In male *Prdm16^csp1/wt^* LV tissue, a similar but less pronounced reduction was detected. Statistical analysis of individual metabolites was performed with non-parametric Wilcoxon Rank Sum test, *indicates p<0.05. **(B)** Univariate analysis reveals in cardiac tissue of *Prdm16^csp1/wt^* mice significant reduction of the amino acid, glycolysis, glycerol, and tricarboxylic acid cycle (TCA) metabolism using combined male and female data. Divergent metabolism of male and female animals was observed for amino acid metabolism, glycolysis, and TCA cycle. **(C)** Lipid analysis was performed with LC-MS using the lipidizer kit (Sciex). Global lipid analysis and evaluation as log2 value of the intensity ratio *Prdm16^csp1/wt^*/controls reveals normal levels for most lipid classes. Strongest regulation was observed for triacylglycerol (TAG) in male *Prdm16^csp1/wt^* hearts. The number (n) of validly detected lipids per class is indicated. **(D)** Selected neutral lipids and sphingolipids critical for the heart, lipids altered in *Prdm16^csp1/wt^* mice, and lipids previously associated with heart function (^†^*Tham* et al. ^22^, ^‡^ *Wittenbecher* et al. ^23^) are presented for *Prdm16^csp1/wt^* cardiac tissue of both sexes and in combination. Values for phospholipids are available in Figure IX in the Data Supplement. **(E)** Signaling activity of the mitogen-activated protein kinase (MAPK) and mTOR pathway was assessed with the Milliplex phosphoprotein magnetic bead system and revealed diminished phosphorylation of insulin receptor (Insr_Tyr1162-1163), phosphatase and tensin homolog (Pten_Ser380), and Akt serine/threonine kinase 2 (Akt_Ser473) in female *Prdm16^csp1/wt^* mice. Statistical analysis of selected lipids was performed with non-parametric Wilcoxon Sum Rank test, * indicates p<0.05. Coloring indicates reduction (red) or increase (blue) of the log2 of ratio by −0.2 or 0.2, respectively.

The adult heart mainly relies on FA oxidation as its primary substrate, so we next explored lipid metabolism with liquid chromatography-mass spectrometry (LC-MS) in depth. Values are presented as log2 from ratio *Prdm16^csp1/wt^*/controls (values >0.2, blue and values <-0.2, red). Evaluation of lipid classes did not reveal significant dysregulation. The strongest alterations were observed in male *Prdm16^csp1/wt^* hearts for triacylglycerol compounds (TAG; increase), dihydroceramides (DCER; reduction), and hexosylceramides (HCER; reduction) (Figure 5C). Currently, there is little known about dysregulation of individual lipid classes in early cardiac dysfunction.^22,23^ Individual lipids are presented as log2 of ratio *Prdm16^csp1/wt^*/controls. Although accumulated lipid classes appear widely normal a minority of individual lipids were dysregulated (Figure IX_A in the Data Supplement). Important FA such as oleic, linoleic, α-linolenic, and arachidonic acid show moderate alteration (Figure 5D). Several diacylglycerols (DAG) appear strongly reduced in *Prdm16^csp1/wt^* hearts for instance DAG(16:1/18:0), which was altered in a study assessing lipid profiles in human heart failure.^23^ DAG(18:1/20:4), which was significantly altered on lipidomic profiling of murine hearts after exercise or pressure overload, was normal in *Prdm16^csp1/wt^* hearts.^22^ Enrichment of individual TAG was significant for TAG47:2-FA18:2, TAG50:4-FA20:4, and TAG51:4- FA18:3 in male and to lesser degree in female *Prdm16^csp1/wt^* cardiac tissue. The sphingolipids sphingomyelin SM(14:0), dihydroceramide DCER(16:0), and ceramide CER(22:1) show strongest reduction in male *Prdm16^csp1/wt^* cardiac tissue. The hexosylceramide, HCER(d18:1/16:0) appeared diminished in hearts of both *Prdm16^csp1/wt^* sexes. Several protective and risk predictive sphingolipids identified by *Wittenbecher et al*. ^23^ were unaffected in *Prdm16^csp1/wt^* mice. The global levels of the cardiac phospholipids phosphatidylcholine (PC) and phosphatidylethanolamine (PE) were unaffected (Figure 5C). Individual PC and PE species appeared consistently enhanced (e.g. PC(18:2/18:3), PE(17:0/22:5), PE(P-18:0/18:0)) or decreased (e.g. PC(18:1/16:1), PE(P-18:1/18:1), PE(P- 18:1/18:2)) in *Prdm16^csp1/wt^* hearts of both sexes (Figure IX_B in the Data Supplement). The level of cardiolipin (CA(18:2(4)), a phospholipid critical for mitochondrial function, appeared moderately elevated in male *Prdm16^csp1/wt^* hearts (Figure 5D). Thus, global lipid metabolism is altered in male *Prdm16^csp1/wt^* hearts with accumulation of TAG.

Cardiac metabolism is controlled by the mitogen-activated protein kinase (MAPK) and mechanistic target of Rapamycin (mTOR) pathways. Using the Milliplex phosphoprotein magnetic bead system we tested the phosphorylation level of key proteins from relevant pathways. Female *Prdm16^csp1/wt^* hearts showed significant inactivation of insulin receptor (Insr_Tyr1162-1163), phosphatase and tensin homolog (Pten_Ser380), and Akt serine/threonine kinase 2 (Akt_Ser473) phosphorylation (Figure 5E). mTOR phosphorylation activation at Mtor_Ser2448 and tuberin (Tsc2_Ser939) showed increased phosphorylation without reaching statistical significance. Thus, diminished phosphorylation of the Insr/Akt pathway activates mTOR signaling and cardiac metabolism adapts accordingly. Altogether, *Prdm16^csp1/wt^* hearts show multiple, substantial metabolic alterations suggesting that monoallelic *Prdm16* inactivation affects cardiac metabolism.

### Tissue energetics and production of protective lipids in Prdm16^csp1/wt^ hearts

To further explore the consequences of the imbalanced cardiac metabolism in *Prdm16^csp1/wt^* mice, we tested the steady-state levels of important cardiac metabolic intermediates, redox molecules, and eicosanoids. Global adenosine triphosphate (ATP) levels were unaffected in *Prdm16^csp1/wt^* cardiac tissue (Figure 6A, Table VIII in the Data Supplement). However, the ratio of adenosine monophosphate to ATP (AMP/ATP) was significantly increased in female *Prdm16^csp1/wt^* hearts. AMP is a critical molecule for sensing metabolic stress conditions and the increased AMP/ATP ratio suggests an abnormal energy state in the *Prdm16^csp1/wt^* cardiac tissue. We also detected increased absolute and relative inosine monophosphate (IMP) values in both male and female *Prdm16^csp1/wt^* hearts (Figure 6B). IMP is the key molecule in purine metabolism and a sensitive measure of the ATP turnover.^24^ The level of hypoxanthine, which serves as precursor and degradation product of IMP, was elevated in male *Prdm16^csp1/wt^* hearts. The ratio of reduced to oxidized nicotinamide adenine dinucleotide (NADH/NAD+), serving as central hydride donor for oxidative phosphorylation and in many other redox reactions^25^, was increased in female *Prdm16^csp1/wt^* hearts (Figure 6C). The ratio of reduced to oxidized glutathione (GSH/GSSG) was diminished in female *Prdm16^csp1/wt^* hearts suggesting accelerated cellular GSH consumption and redox imbalance (Figure 6D). The levels of creatine, phosphor-creatine/creatine, and acetyl-CoA appeared normal. The level of 4-hydroxynonenal (4-HNE), a marker of lipid peroxidation, was unaffected. These findings suggest accelerated ATP turnover and oxidative stress but no lipid peroxidation on *Prdm16* inactivation in particular in female *Prdm16^csp1/wt^* hearts.

**Figure 6.**
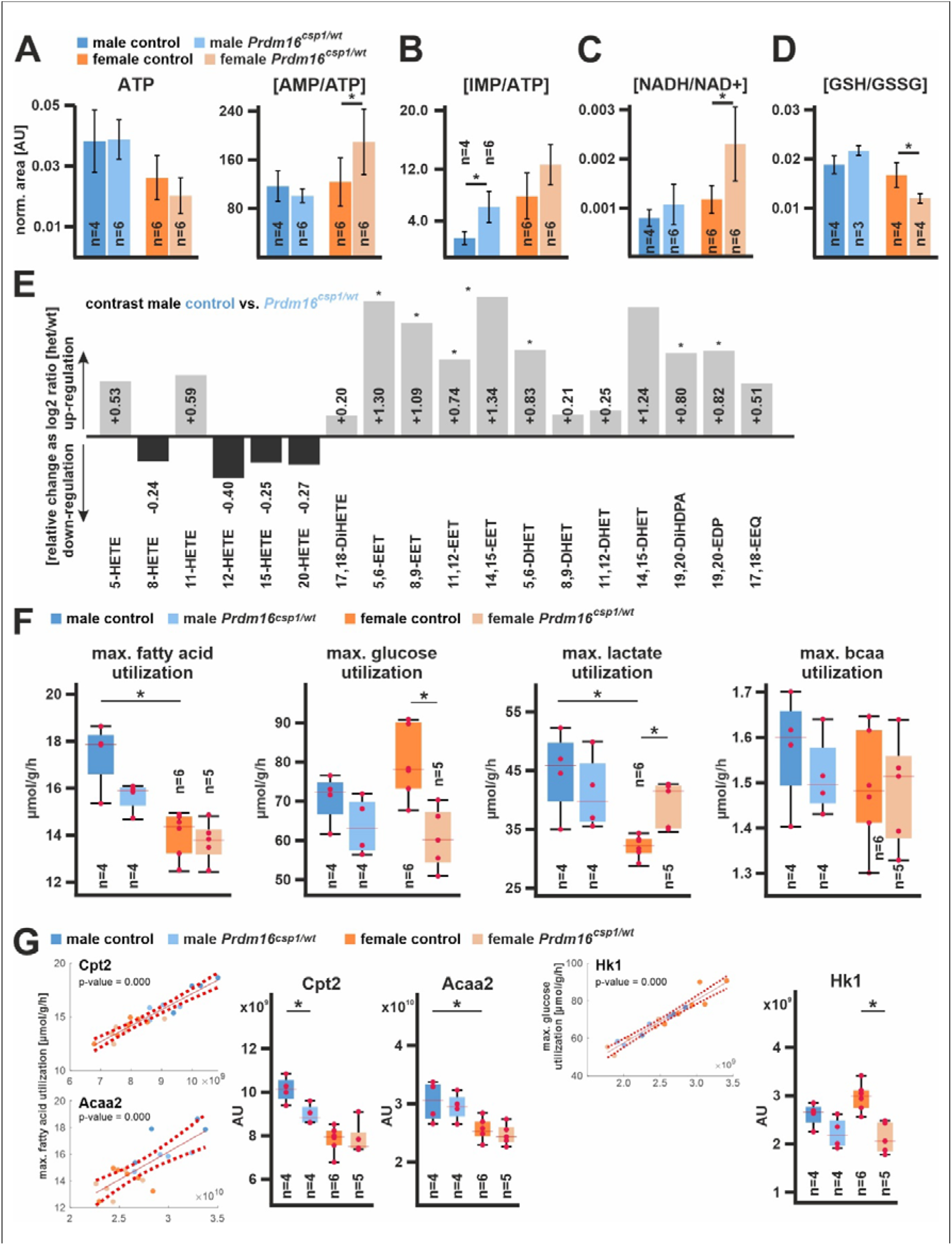
Nutrient metabolism in *Prdm16^csp1/wt^* cardiac tissue. **(A)** Assessment of metabolites critical for energy metabolism using LV tissue and LC-MS. Normalized values are shown for adenosine triphosphate (ATP) and the ratio of adenosine monophosphate (AMP) vs. ATP (AMP/ATP). **(B)** The ratio of inosine monophosphate (IMP) vs. ATP (IMP/ATP) is increased in *Prdm16^csp1/wt^* hearts. **(C)** The ratio of reduced vs. oxidized nicotinamide adenine dinucleotide (NADH/NAD+) is increased in female *Prdm16^csp1/wt^* hearts. **(D)** Oxidative capacity of cardiac tissues was assessed with the ratio of reduced vs. oxidized glutathione (GSH/GSSG). Female *Prdm16^csp1/wt^* hearts show a significantly reduced GSH/GSSG ratio. **(E)** Eicosanoids were measured in male *Prdm16^csp1/wt^* hearts with LC/ESI-MS-MS. Data are presented as log2 *Prdm16^csp1/wt^*/controls ratio with down- or up-regulation as black or grey bars, respectively. All epoxyeicosatrienoic (EET) and dihydroxyeicosatrienoic (DHET) acids are increased in *Prdm16^csp1/wt^* cardiac tissue. Corresponding absolute measurements are available in Table VIII in the Data Supplement. **(F)** Modelling of major cardiac metabolic processes occurred with CARDIOKIN1 ^46^ using protein expression data. Differences in maximal substrate utilization for fatty acids (FA), glucose, lactate, and branched chain amino acids (bcaa). Box plots show median and 25% quartile. Red dots depict maximal capacities for individual animals. **(G)** Individual protein impact for FA and glucose metabolism was correlated for carnitine palmitoyltransferase 2 (Cpt2), acetyl-CoA acyltransferase 2 (Acaa2), and hexokinase 1 (Hk1) using linear regression analysis of the maximal substrate utilization vs. protein abundance (red dashed line indicates confidence interval, 95%). Box plots show median and 25% quartile. Red dots depict Cpt2, Acaa2, and Hk1 maximal capacities for individual animals. Statistical analysis of individual metabolites and processes was performed with unpaired t-test, * indicates p<0.05.

Apart from their function as nutritional source FA serve as reactant for biosynthesis of hydroxyeicosatetraenoic acids (HETE) and epoxyeicosatrienoic acids (EET)^26,27^. All HETE species, which are synthesized from arachidonic acid, FA(20:4) by lipoxygenases (LOX), were normal in male *Prdm16^csp1/wt^* cardiac tissue (Figure 6E, Table IX in the Data Supplement). 20-HETE, which is synthesized from FA(20:4) by cytochrome P450 (CYP), was also unaffected. All tested individual EET, which originate from CYP activity, and summed EET were significantly increased. Consistently, also the water soluble dihydroxyeicosatrienoic acids (DHET), representing the corresponding EET hydrolyzation products, were significantly increased. The anti-inflammatory 19,20-epoxydocosapentaenoic acid (19,20-EDP) and 17,18-epoxyeicosatetraenoic acid (17,18-EEQ) were also significantly increased. These observations suggest activation of CYP mediated bioactive, lipid production (EETs and 19,20-EDP) in male *Prdm16^csp1/wt^* cardiac tissue.

To assess the global metabolism of *Prdm16^csp1/wt^* cardiac tissue, we modelled major metabolic pathways with CARDIOKIN1 *in silico*. As input data we used quantitative protein expression data of metabolically important proteins measured with LC-MS/MS (Figure 4). The calculated maximal utilization rates of the given metabolic condition are upper estimates that will probably not be reached under physiological conditions. However, these maximal utilization rates allow estimation of the physiological capacity. The maximal FA and glucose utilization rates were different between cardiac the tissue of male and female control hearts. Females had lower FA but higher glucose utilization capacity compared to male controls (Figure 6F). In addition, the maximal lactate utilization rate was lower in female controls. In male *Prdm16^csp1/wt^* cardiac tissue the maximal FA utilization was decreased but unaffected in females. In contrast, female *Prdm16^csp1/wt^* cardiac tissue had a diminished maximal glucose utilization, while the males were unaffected. Maximal lactate utilization was increased in female *Prdm16^csp1/wt^* cardiac tissue. Maximal ATP production and O_2_ consumption were modelled under fasting and postprandial conditions. Under both conditions the cardiac tissue of female controls showed diminished maximal ATP production and O_2_ consumption compared to male controls (Figure X in the Data Supplement). The maximal utilization rate gives a net measure involving several members of the given metabolic pathway. Thus, we aimed to identify enzymes critical for FA and glucose metabolism in *Prdm16^csp1/wt^* cardiac tissue. For this purpose, the metabolic utilization and protein abundance was correlated for each protein and animal. The strongest correlations, indicated by low p-values, were identified for carnitine palmitoyltransferase 2 (Cpt2), acetyl-CoA acyltransferase 2 (Acaa2), hexokinase 1 (Hk1), and others (Figure 6G, Figure XI in the Data Supplement). In controls, Cpt2 and Acaa2, two determinants of the mitochondrial beta-oxidation activity, showed lower maximal capacity in females supporting their diminished FA metabolism (Figure 6G, Figure XII in the Data Supplement). Lower Cpt2 capacity in male *Prdm16^csp1/wt^* cardiac tissue partly explains diminished male FA metabolism. Consistent with protein expression data, female and male *Prdm16^csp1/wt^* cardiac tissue showed diminished Hk1 utilization. Hk1 mediates phosphorylation of D-glucose to D-glucose 6-phosphate and represents the initial step of glycolysis. Altogether, these findings support imbalanced substrate metabolism and oxidative stress on monoallelic *Prdm16* inactivation.

## Discussion

Our study suggests that metabolic dysregulation is an early event in *Prdm16* associated cardiac dysfunction and precedes substantial transcriptional dysregulation. Metabolic changes upon *Prdm16* inactivation appear to contribute to induce early and late-stage cardiac pathologies, which may result in cardiac hypoplasia^9^ and hypertrophy^28^, respectively. Pyroxd2 and Pbxip1 are novel modulators of cardiac function, reflecting the central role of metabolism in the *Prdm16* associated cardiac phenotype. Moreover, our study detects a more pronounced molecular and structural phenotype in *Prdm16^csp1/wt^* females than males, with sex having a larger effect than the *Prdm16* genotype. The differential response in maximal FA and glucose utilization of male and female *Prdm16^csp1/wt^* hearts suggest a diminished capacity of females to scope with metabolic challenges.

### PRDM16 cardiomyopathy

Mutation of *PRDM16* is a cause of cardiomyopathy associated with DCM and LVNC.^4^ More recently, truncating variants in *PRDM16* were identified by *in silico* analysis as one of three specific LVNC variant classes.^8^ These genetic observations and the *Prdm16^csp1/wt^* phenotype reinforce the human genetic case that heterozygous *PRDM16* inactivation/truncation is sufficient to cause cardiomyopathy. Functional evidence from *Prdm16^csp1/wt^* mice, mirroring the genotype of patients with *PRDM16* mutation, will increase the ClinGen evidence from *limited* to *moderate* for the association of *PRDM16* with cardiomyopathy (www.clinicalgenome.org).

In mice, homozygous *Prdm16* inactivation induces discordant phenotypes in different genetic backgrounds resulting in either early postnatal lethality (*Xmlc2Cre;Prdm16^flox/flox^*) or cardiac dysfunction with fibrosis at adult stages (*Mesp1Cre;Prdm16^flox/flox^*, *Myh6Cre;Prdm16^flox/flox^*).^10,28,29^ *Xmlc2Cre* and *Mesp1Cre* strains are expected to inactivate *Prdm16* upon very early heart development.^30^ Discrepancies in penetrance of the phenotypes could be attributed to incomplete spatial Cre-mediated *Prdm16* inactivation and residual Prdm16 levels in early cardiomyocyte progenitor cells, endothelia, or cardiac fibroblasts. Consistent with neuronal studies, it is likely that *Prdm16* has additional effects on cardiac stem and progenitor cell function compared with differentiated, adult cardiomyocytes.^31^ Indeed, adult *Prdm16* inactivation using a tamoxifen-inducible mouse model (*αMHC-MerCreMer;Prdm16^flox/flox^*) resulted in viable mice without an overt cardiac phenotype.^10^ Homozygous *PRDM16* mutation has not been identified in patients so far but based on animal studies^10,28,29^, biallelic *PRDM16* inactivation likely induces a highly penetrant, severe cardiac phenotype in humans that may not survive to birth. In *Prdm16^csp1/wt^* hearts we did not observe alterations in myocardial compaction or alterations of the gene circuit involving Tbx5 and Hand1.^10^ This is possibly due to the heterozygous nature of the *Prdm16* inactivation in our model. Another important feature is that *Prdm16^csp1/wt^* hearts do not show perturbation of the sarcomere at either a structural or a molecular level. This suggests that contractile dysfunction in the *PRDM16* cardiomyopathy originates from other mechanisms such as energy restriction or metabolic stress affecting contractility and/or ion homeostasis.

### Prdm16 differentially compromises metabolism in early and late stage cardiac pathology

Studies from adipose tissue show that PRDM16 orchestrates adipocyte differentiation via interaction with proteins such as PPARA, PPARG, MED1, CEBPD, peroxisome proliferator-activated receptor gamma coactivator 1-alpha, beta (PPARGC1A, PPARGC1B), or uncoupling protein 1 (UCP1).^3,12,13,15–17,32,33^ Consequently, we explored a selection of validated PRDM16 targets in *Prdm16^csp1/wt^* hearts and in a transcriptomic screen of human adult DCM (Table VII in the Data Supplement).^18^ We found only three PRDM16 targets, namely PPARA, MED1, and CEBPD dysregulated in this DCM transcriptomic screen. Moreover, the transcriptional targets Tbx5, Hand1 identified in *Xmlc2Cre;Prdm16^flox/flox^* hearts appeared not be regulated in *Prdm16^csp1/wt^* hearts.^10^ This suggests: *i*) in cardiomyocytes PRDM16 steers diverse regulatory programs compared to other differentiated cell types, *ii*) expression changes are time sensitive and depend on the progenitor or differentiation stage, and *iii*) early molecular changes or adaptations in PRDM16 cardiac pathology are not associated with broad, significant expression changes. In the absence of major transcriptional effects in our model, we investigated possible metabolic alterations.

The initial evidence that metabolic alteration may be associated with the cardiac *PRDM16* phenotype came from a study analyzing homozygous *Mesp1Cre;Prdm16^flox/flox^* mice. These mice develop cardiac hypertrophy, diminished heart function, fibrosis, reduced mitochondrial content, and diminished acylcarnitine levels at 12 months of age.^28^ *Mesp1Cre;Prdm16^flox/flox^* hearts show upregulation of transcripts involved in glucose metabolism, transcriptional downregulation of lipid metabolism, and oxidative stress gene expression. Metabolic high-fat diet challenge of young *Mesp1Cre;Prdm16^flox/flox^* mice induced cardiac dysfunction as well as hypertrophy already at 3 months of age compared to a later phenotype onset at 9 months without high-fat diet. This suggests that complete *Prdm16* inactivation in the heart leads to diminished mitochondrial volume and sensitivity to substrate availability ultimately resulting in cardiomyocyte hypertrophy by activating distinctive gene programs.^28^ Exploration of the heart phenotype in *Xmlc2Cre;Prdm16^flox/flox^* mice also detected downregulation of transcripts associated with mitochondrial biogenesis/function and FA metabolism.^10^ Together, both models analyzing homozygous *Prdm16* inactivation demonstrate cardiac phenotypes that are associated with diminished metabolic capacity eventually leading to cardiac growth and hypertrophy. In contrast, heterozygous *Prdm16^csp1/wt^* hearts are hypoplastic with no evidence of mitochondrial volume changes, demonstrating that metabolic alterations are early events in *Prdm16* inactivation.

A differential response in maximal FA or glucose utilization of male and female *Prdm16^csp1/wt^* is supported by sex-specific transcriptional dysregulation. In male *Prdm16^csp1/wt^* hearts the strongest up- and downregulation was observed for *Ubiad1* and hemoglobin’s (*Hba-a1, Hbb-bs*), respectively. Ubiad1 is a prenyltransferase that is involved in ubiquinone (CoQ10) synthesis, thus, increasing redox tolerance and providing cardiovascular protection.^34^ Cellular downregulation of Hba-a1 and Hbb-ba diminishes oxygen supply and may blunt cardiac oxidative stress in the setting of restricted FA oxidation. Female *Prdm16^csp1/wt^* hearts show strongest up- and downregulation of the metabolism associated transcripts *Nr1d1* and *Aldh3a1*, respectively. NR1D1 is a ligand-regulated transcriptional repressor impacting metabolic regulation, cellular differentiation, or circadian rhythm control.^35^ Upon adipogenesis, *Nr1d1* gene silencing affects brown adipocyte differentiation and attenuates *Prdm16* expression suggesting an interaction of both.^35^ A recent study exploring the impact of shift work on cardiac reperfusion injury demonstrated diminished myocardial Nr1d1 expression suggesting a more general role of Nr1d1 in cardiomyocyte function.^36^ Aldh3a1 is involved in oxidation and detoxification of lipid peroxids. Genetic inactivation of *Aldh3a1* in zebrafish increases 4-HNE levels and impairs glucose homeostasis.^37^ Thus, Aldh3a1 reduction is likely associated with metabolic adaptation of female *Prdm16^csp1/wt^* glucose utilization. Overall, early transcriptional changes in *Prdm16^csp1/wt^* hearts are distinct from those observed in later stage pathology.

### Pyroxd2 and Pbxip1 are novel modulators of cardiac function

In line with a central role of metabolism for PRDM16-associated cardiac phenotypes, we found concordant upregulation of Pyroxd2 and Pbxip1 both of which have been implicated in the regulation of energy metabolism. PBXIP1 interacts with several proteins such as the transcription factor Pre-B cell leukemia factor 1 (PBX1)^38^, microtubules^38^, estrogen receptors 1 and 2 (ESR1, ESR2)^39^, and AMP-activated protein kinase (AMPK).^40^ Moreover, PBXIP1 regulates the activity of MAPK and mTOR signaling.^21^ The function of Pbxip1 in the heart is largely unknown. In a prior genetic screen, the inhibition of MAPK signaling pathway proteins increased PBXIP1 phosphorylation.^41^ In contrast, activation of the MAPK signaling cascade increased PBXIP1 protein expression and murine PBXIP1 overexpression stimulated cardiac hypertrophy.^41^ In *Prdm16^csp1/wt^* hearts we found Pbxip1 upregulation associated with diminished MAPK signaling activity. Mutation of *PBXIP1* has not been associated with human heart disease; however, mutation of its interacting transcription factor PBX1 has been linked to syndromic congenital heart defects.^42^ An interesting aspect of PBXIP1 is its interaction and activation of ESR1 and ESR2, a potential mechanisms of sex specific effects.^39^ Our data implicate PBXIP1 in cardiac growth, metabolism, and hypertrophy which await further characterization under healthy and diseased conditions. In the context of *Prdm16* heterozygous mutants, Pbxip1 may orchestrate the adaptive responses of important signaling cascades like AMPK, mTOR, or MAPK.

The cardiac function of Pyroxd2 is unknown. PYROXD2 is an oxidoreductase of the inner mitochondrial membrane/matrix that interacts with mitochondrial complex IV.^20^ Genetic inactivation of PYROXD2 in hepatic cell lines decreased the mitochondrial membrane potential, complex IV activity, ATP content, and mitochondrial DNA copy number.^20^ PYROXD2 inactivation also increased mitochondrial reactive oxygen species and the number of immature mitochondria.^20^ Compound heterozygous genetic variants in *PYROXD2* were detected in a single patient with a severe infantile metabolic disorder.^43^ Molecular workup in patient fibroblasts demonstrated a mito-ribosomal defect characterized by increased mitochondrial superoxide levels, elevated sensitivity to metabolic stress, decreased complex I subunit proteins, and diminished mitochondrial ribosome levels.^43^ Mutation of a related human oxidoreductase, *PYROXD1*, results in early-onset skeletal myopathy.^44^ Another study, investigating molecular signatures in skeletal muscle from heart failure patients, detected upregulation of *PYROXD2* transcripts.^45^ These data suggest that Pyroxd2 upregulation in *Prdm16^csp1/wt^* hearts may compensate for metabolic and oxidative stress conditions.

### Sex specific aspects of Prdm16 inactivation in cardiac metabolism

Our study detects a more pronounced cardiac and molecular phenotype in *Prdm16^csp1/wt^* females compared to males. The original cytogenetic association of *PRDM16* inactivation with cardiomyopathy was described in 18 patients with 1p36 syndrome.^4^ Among these, 16 individuals were females and only two were males.^4^ This may point to a sex specific penetrance of the *PRDM16* cardiomyopathy. Apparently, female *Prdm16^csp1/wt^* mice compensate the cardiac metabolic disturbance less efficiently than males. Metabolic modelling with CARDIOKIN1 revealed diminished FA, increased glucose, and reduced lactate utilization in the hearts of female compared with male *Prdm16^wt/wt^* mice.^46^ This establishes clear sexual dimorphism for cardiac metabolism under basal healthy conditions.^47^ Under normal conditions the heart generates ATP mainly via FA oxidation and glucose utilization (glycolysis, pyruvate supply to TCA).^48^ Female *Prdm16^csp1/wt^* hearts show diminished maximal glucose and unaffected FA utilization. The increase in the maximal lactate utilization capacity in female *Prdm16^csp1/wt^* hearts likely reflects an adaptation to metabolic stress conditions. Male *Prdm16^csp1/wt^* hearts show diminished maximal FA utilization, which is in line with the TAG accumulation we observed. Male *Prdm16^csp1/wt^* hearts expose elevated EET levels, which is in line with a recent study demonstrating increased EET levels/turnover in LV biopsies from DCM patients.^49^ This supports distinct mechanisms of lipid metabolism in male hearts. Whether hormonal differences are the cause of systemically reduced fat content and the more advanced pathology in female *Prdm16^csp1/wt^* mice is unclear. The known association of Pbxip1 and Esr1/Esr2 activity as well as Nr1d1 provides a potential explanation for the sexual dimorphism in the regulation of substrate utilization. Transcript as well as protein level of *Prdm16^csp1/wt^* hearts demonstrate significant sex effects. Indeed, sex has a larger effect on metabolic parameters than the *Prdm16* genotype at least during this investigated early pathological window. Our findings suggest a sexual dimorphism for the *PRDM16* associated cardiomyopathy with probably earlier, more penetrant phenotype expression in females.

**Figure 7.**
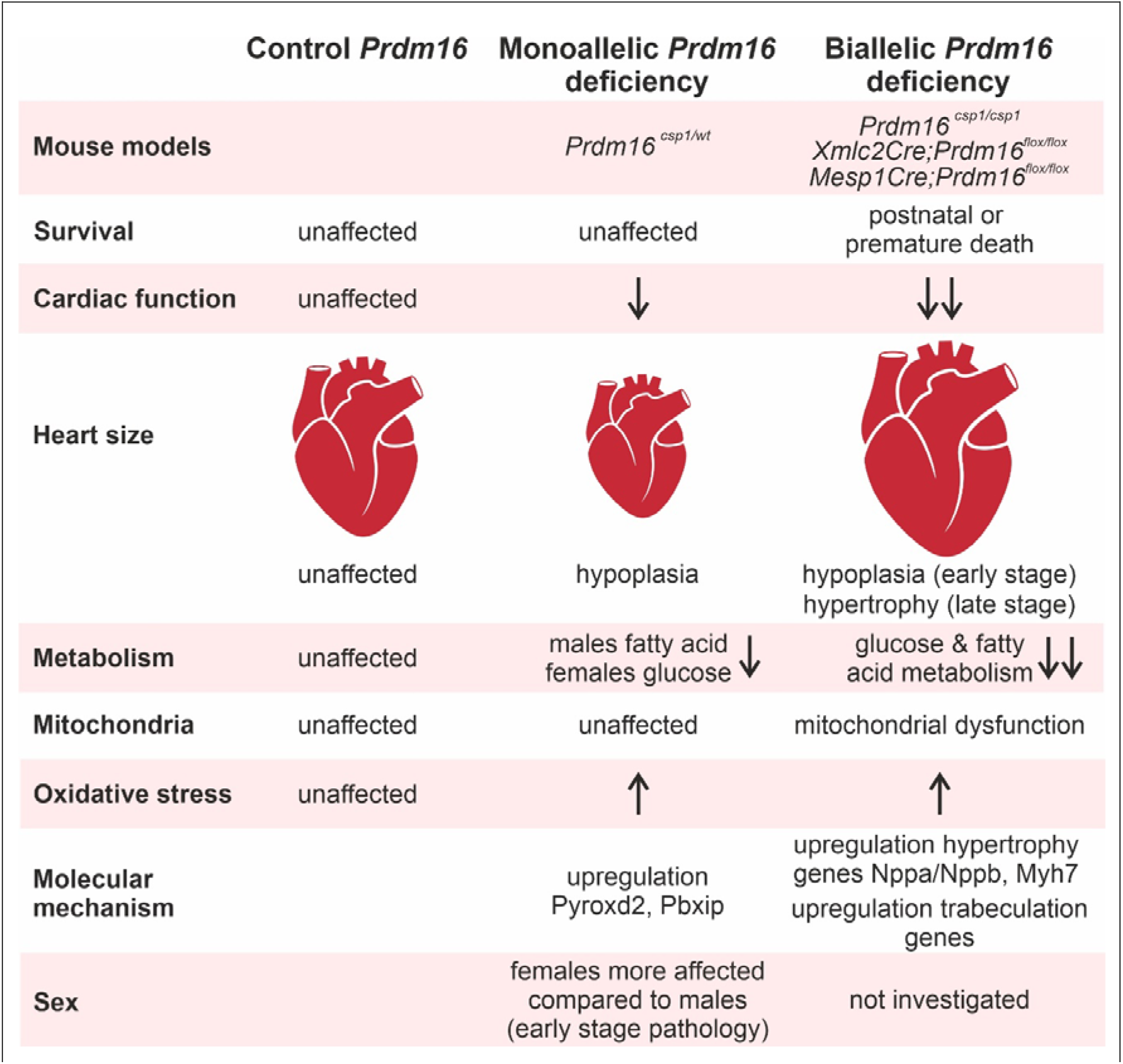
Prdm16 is an early regulator of cardiac metabolism. Assessing different mouse models for the *PRDM16* associated cardiomyopathy reveals distinct cardiac phenotypes after mono- or biallelic Prdm16 inactivation. Metabolism is differentially affected at early (*Prdm16^csp1/wt^*) and late pathology stage (*Xmlc2Cre;Prdm16^flox/flox^, Mesp1Cre;Prdm16^flox/flox^*).^10,28^

## Nonstandard Abbreviation and Acronyms

DCM: dilated cardiomyopathy
LVNC: left ventricular noncompaction cardiomyopathy
Pbxip1: pre B-cell leukemia transcription factor interacting protein 1
*PRDM16*: human PR/SET domain 16 gene
PRDM16: human PR/SET domain 16 protein
*Prdm16*: mouse PR/SET domain 16 gene
Prdm16: mouse PR/SET domain 16 protein
*Prdm16^csp1/wt^*: mice carrying the csp1 mutant allele heterozygous
Pyroxd2: pyridine nucleotide-disulphide oxidoreductase domain 2

## Study Limitations

Female *Prdm16^csp1/wt^* mice have diminished relative total body fat content and increased relative muscle content. This mirrors either the female tissue specific metabolism and/or points towards a systemic factor affecting tissue development. Identification of such factor would be important to understand the systemic (endocrine) role of Prdm16 and to test therapeutic approaches targeting adipose tissue. Our study does not assess early metabolic changes in cardiac progenitor cells. Analysis of *Prdm16^csp1/wt^* hearts with *in silico* tools and individual metabolite measurements suggest an accelerated metabolism. However, we did not perform kinetic experiments to demonstrate accelerated turnover of selected metabolic pathways.

## Acknowledgments

We thank the Berlin Institute of Health (BIH), Core Facility Genomics, Berlin, Germany for providing the high throughput sequencing platform (Tatjana Borodina). The MDC, animal phenotyping facility performed mouse physiological experiments (Stefanie Schelenz, Martin Taube). We thank the Advanced Light Microscopy Technology Platform of the Max-Delbrück-Center for Molecular Medicine, Berlin for the general and technical support (Anca Margineanu, Anje Sporbert). We thank the Charité electron microscopy facility (Petra Schrade, Sara Timm, Matthias Ochs). We thank the MS Omics for metabolic analysis (Lea Johnsen, Morten Danielsen).

## References

1. Burke MA, Cook SA, Seidman JG, Seidman CE. Clinical and Mechanistic Insights Into the Genetics of Cardiomyopathy. Journal of the American College of Cardiology. 2016;68:2871–2886. doi: 10.1016/j.jacc.2016.08.079

2. Kuhnisch J, Herbst C, Al-Wakeel-Marquard N, Dartsch J, Holtgrewe M, Baban A, Mearini G, Hardt J, Kolokotronis K, Gerull B, et al. Targeted panel sequencing in pediatric primary cardiomyopathy supports a critical role of TNNI3. Clinical genetics. 2019;96:549–559. doi: 10.1111/cge.13645

3. Ishibashi J, Seale P. Functions of Prdm16 in thermogenic fat cells. Temperature. 2015;2:65–72.

4. Arndt AK, Schafer S, Drenckhahn JD, Sabeh MK, Plovie ER, Caliebe A, Klopocki E, Musso G, Werdich AA, Kalwa H, et al. Fine mapping of the 1p36 deletion syndrome identifies mutation of PRDM16 as a cause of cardiomyopathy. American journal of human genetics. 2013;93:67–77. doi: 10.1016/j.ajhg.2013.05.015

5. Delplancq G, Tarris G, Vitobello A, Nambot S, Sorlin A, Philippe C, Carmignac V, Duffourd Y, Denis C, Eicher JC, et al. Cardiomyopathy due to PRDM16 mutation: First description of a fetal presentation, with possible modifier genes. American journal of medical genetics Part C, Seminars in medical genetics. 2020;184:129–135. doi: 10.1002/ajmg.c.31766

6. van Waning JI, Caliskan K, Hoedemaekers YM, van Spaendonck-Zwarts KY, Baas AF, Boekholdt SM, van Melle JP, Teske AJ, Asselbergs FW, Backx A, et al. Genetics, Clinical Features, and Long-Term Outcome of Noncompaction Cardiomyopathy. Journal of the American College of Cardiology. 2018;71:711–722. doi: 10.1016/j.jacc.2017.12.019

7. Long PA, Evans JM, Olson TM. Diagnostic Yield of Whole Exome Sequencing in Pediatric Dilated Cardiomyopathy. J Cardiovasc Dev Dis. 2017;4. doi: 10.3390/jcdd4030011

8. Mazzarotto F, Hawley MH, Beltrami M, Beekman L, de Marvao A, McGurk KA, Statton B, Boschi B, Girolami F, Roberts AM, et al. Systematic large-scale assessment of the genetic architecture of left ventricular noncompaction reveals diverse etiologies. Genet Med.2021;23:856–864. doi: 10.1038/s41436-020-01049-x

9. Bjork BC, Turbe-Doan A, Prysak M, Herron BJ, Beier DR. Prdm16 is required for normal palatogenesis in mice. Human molecular genetics. 2010; 19:774–789. doi: 10.1093/hmg/ddp543

10. Wu T, Liang Z, Zhang Z, Liu C, Zhang L, Gu Y, Peterson KL, Evans SM, Fu XD, Chen J. PRDM16 Is a Compact Myocardium-Enriched Transcription Factor Required to Maintain Compact Myocardial Cardiomyocyte Identity in Left Ventricle. Circulation. 2022;145:586–602. doi: 10.1161/CIRCULATIONAHA.121.056666

11. Jiang N, Yang M, Han Y, Zhao H, Sun L. PRDM16 Regulating Adipocyte Transformation and Thermogenesis: A Promising Therapeutic Target for Obesity and Diabetes. Frontiers in pharmacology. 2022;13:870250. doi: 10.3389/fphar.2022.870250

12. Kajimura S, Seale P, Kubota K, Lunsford E, Frangioni JV, Gygi SP, Spiegelman BM. Initiation of myoblast to brown fat switch by a PRDM16-C/EBP-beta transcriptional complex. Nature.2009;460:1154–1158. doi: 10.1038/nature08262

13. Kajimura S, Seale P, Tomaru T, Erdjument-Bromage H, Cooper MP, Ruas JL, Chin S, Tempst P, Lazar MA, Spiegelman BM. Regulation of the brown and white fat gene programs through a PRDM16/CtBP transcriptional complex. Genes & development. 2008;22:1397–1409. doi: 10.1101/gad.1666108

14. Seale P, Kajimura S, Spiegelman BM. Transcriptional control of brown adipocyte development and physiological function--of mice and men. Genes & development. 2009;23:788–797. doi: 10.1101/gad.1779209

15. Seale P, Kajimura S, Yang W, Chin S, Rohas LM, Uldry M, Tavernier G, Langin D, Spiegelman BM. Transcriptional control of brown fat determination by PRDM16. Cell metabolism. 2007;6:38–54. doi: 10.1016/j.cmet.2007.06.001

16. Harms MJ, Lim HW, Ho Y, Shapira SN, Ishibashi J, Rajakumari S, Steger DJ, Lazar MA, Won KJ, Seale P. PRDM16 binds MED1 and controls chromatin architecture to determine a brown fat transcriptional program. Genes & development. 2015;29:298–307. doi: 10.1101/gad.252734.114

17. Seale P, Bjork B, Yang W, Kajimura S, Chin S, Kuang S, Scime A, Devarakonda S, Conroe HM, Erdjument-Bromage H, et al. PRDM16 controls a brown fat/skeletal muscle switch. Nature. 2008;454:961–967. doi: 10.1038/nature07182

18. Heinig M, Adriaens ME, Schafer S, van Deutekom HWM, Lodder EM, Ware JS, Schneider V, Felkin LE, Creemers EE, Meder B, et al. Natural genetic variation of the cardiac transcriptome in non-diseased donors and patients with dilated cardiomyopathy. Genome biology.2017;18:170. doi: 10.1186/s13059-017-1286-z

19. Pomaznoy M, Ha B, Peters B. GOnet: a tool for interactive Gene Ontology analysis. BMC Bioinformatics. 2018;19:470. doi: 10.1186/s12859-018-2533-3

20. Wang T, Xie X, Liu H, Chen F, Du J, Wang X, Jiang X, Yu F, Fan H. Pyridine nucleotide-disulphide oxidoreductase domain 2 (PYROXD2): Role in mitochondrial function. Mitochondrion. 2019;47:114–124. doi: 10.1016/j.mito.2019.05.007

21. Khumukcham SS, Manavathi B. Two decades of a protooncogene HPIP/PBXIP1: Uncovering the tale from germ cell to cancer. Biochim Biophys Acta Rev Cancer. 2021;1876:188576. doi: 10.1016/j.bbcan.2021.188576

22. Tham YK, Bernardo BC, Huynh K, Ooi JYY, Gao XM, Kiriazis H, Giles C, Meikle PJ, McMullen JR. Lipidomic Profiles of the Heart and Circulation in Response to Exercise versus Cardiac Pathology: A Resource of Potential Biomarkers and Drug Targets. Cell reports. 2018;24:2757–2772. doi: 10.1016/j.celrep.2018.08.017

23. Wittenbecher C, Eichelmann F, Toledo E, Guasch-Ferre M, Ruiz-Canela M, Li J, Aros F, Lee CH, Liang L, Salas-Salvado J, et al. Lipid Profiles and Heart Failure Risk: Results From Two Prospective Studies. Circulation research. 2021;128:309–320. doi: 10.1161/CIRCRESAHA.120.317883

24. Johnson TA, Jinnah HA, Kamatani N. Shortage of Cellular ATP as a Cause of Diseases and Strategies to Enhance ATP. Frontiers in pharmacology. 2019; 10:98. doi: 10.3389/fphar.2019.00098

25. Xie N, Zhang L, Gao W, Huang C, Huber PE, Zhou X, Li C, Shen G, Zou B. NAD(+) metabolism: pathophysiologic mechanisms and therapeutic potential. Signal Transduct Target Ther. 2020;5:227. doi: 10.1038/s41392-020-00311-7

26. Schunck WH, Konkel A, Fischer R, Weylandt KH. Therapeutic potential of omega-3 fatty acid-derived epoxyeicosanoids in cardiovascular and inflammatory diseases. Pharmacology & therapeutics. 2018;183:177–204. doi: 10.1016/j.pharmthera.2017.10.016

27. Wang B, Wu L, Chen J, Dong L, Chen C, Wen Z, Hu J, Fleming I, Wang DW. Metabolism pathways of arachidonic acids: mechanisms and potential therapeutic targets. Signal Transduct Target Ther. 2021;6:94. doi: 10.1038/s41392-020-00443-w

28. Cibi DM, Bi-Lin KW, Shekeran SG, Sandireddy R, Tee N, Singh A, Wu Y, Srinivasan DK, Kovalik JP, Ghosh S, et al. Prdm16 Deficiency Leads to Age-Dependent Cardiac Hypertrophy, Adverse Remodeling, Mitochondrial Dysfunction, and Heart Failure. Cell reports.2020;33:108288. doi: 10.1016/j.celrep.2020.108288

29. Nam JM, Lim JE, Ha TW, Oh B, Kang JO. Cardiac-specific inactivation of Prdm16 effects cardiac conduction abnormalities and cardiomyopathy-associated phenotypes. American journal of physiology Heart and circulatory physiology. 2020;318:H764–H777. doi: 10.1152/ajpheart.00647.2019

30. Bruneau BG. Transcriptional regulation of vertebrate cardiac morphogenesis. Circulation research. 2002;90:509–519. doi: 10.1161/01.res.0000013072.51957.b7

31. Leszczynski P, Smiech M, Parvanov E, Watanabe C, Mizutani KI, Taniguchi H. Emerging Roles of PRDM Factors in Stem Cells and Neuronal System: Cofactor Dependent Regulation of PRDM3/16 and FOG1/2 (Novel PRDM Factors). Cells. 2020;9. doi: 10.3390/cells9122603

32. Iida S, Chen W, Nakadai T, Ohkuma Y, Roeder RG. PRDM16 enhances nuclear receptor-dependent transcription of the brown fat-specific Ucp1 gene through interactions with Mediator subunit MED1. Genes & development. 2015;29:308–321. doi: 10.1101/gad.252809.114

33. Seale P, Conroe HM, Estall J, Kajimura S, Frontini A, Ishibashi J, Cohen P, Cinti S, Spiegelman BM. Prdm16 determines the thermogenic program of subcutaneous white adipose tissue in mice. The Journal of clinical investigation. 2011;121:96–105. doi: 10.1172/JCI44271

34. Mugoni V, Postel R, Catanzaro V, De Luca E, Turco E, Digilio G, Silengo L, Murphy MP, Medana C, Stainier DY, et al. Ubiad1 is an antioxidant enzyme that regulates eNOS activity by CoQ10 synthesis. Cell. 2013;152:504–518. doi: 10.1016/j.cell.2013.01.013

35. Nam D, Chatterjee S, Yin H, Liu R, Lee J, Yechoor VK, Ma K. Novel Function of Rev-erbalpha in Promoting Brown Adipogenesis. Scientific reports. 2015;5:11239. doi: 10.1038/srep11239

36. Zhao Y, Lu X, Wan F, Gao L, Lin N, He J, Wei L, Dong J, Qin Z, Zhong F, et al. Disruption of Circadian Rhythms by Shift Work Exacerbates Reperfusion Injury in Myocardial Infarction. Journal of the American College of Cardiology. 2022;79:2097–2115. doi: 10.1016/j.jacc.2022.03.370

37. Lou B, Boger M, Bennewitz K, Sticht C, Kopf S, Morgenstern J, Fleming T, Hell R, Yuan Z, Nawroth PP, et al. Elevated 4-hydroxynonenal induces hyperglycaemia via Aldh3a1 loss in zebrafish and associates with diabetes progression in humans. Redox biology.2020;37:101723. doi: 10.1016/j.redox.2020.101723

38. Abramovich C, Shen WF, Pineault N, Imren S, Montpetit B, Largman C, Humphries RK. Functional cloning and characterization of a novel nonhomeodomain protein that inhibits the binding of PBX1-HOX complexes to DNA. The Journal of biological chemistry.2000;275:26172–26177. doi: 10.1074/jbc.M001323200

39. Wang X, Yang Z, Zhang H, Ding L, Li X, Zhu C, Zheng Y, Ye Q. The estrogen receptor-interacting protein HPIP increases estrogen-responsive gene expression through activation of MAPK and AKT. Biochimica et biophysica acta. 2008;1783:1220–1228. doi: 10.1016/j.bbamcr.2008.01.026

40. Moreno D, Viana R, Sanz P. Two-hybrid analysis identifies PSMD11, a non-ATPase subunit of the proteasome, as a novel interaction partner of AMP-activated protein kinase. The international journal of biochemistry & cell biology. 2009;41:2431–2439. doi: 10.1016/j.biocel.2009.07.002

41. Grimes KA, Pyo A, Molkentin JD. Abstract 918: Pre-B-cell Leukemia Homeobox Interacting Protein 1 is a Novel Regulator of Growth Signaling in the Heart. Circulation research.2019;125.

42. Alankarage D, Szot JO, Pachter N, Slavotinek A, Selleri L, Shieh JT, Winlaw D, Giannoulatou E, Chapman G, Dunwoodie SL. Functional characterization of a novel PBX1 de novo missense variant identified in a patient with syndromic congenital heart disease. Human molecular genetics. 2020;29:1068–1082. doi: 10.1093/hmg/ddz231

43. Van Bergen NJ, Hock DH, Spencer L, Massey S, Stait T, Stark Z, Lunke S, Roesley A, Peters H, Lee JY, et al. Biallelic Variants in PYROXD2 Cause a Severe Infantile Metabolic Disorder Affecting Mitochondrial Function. International journal of molecular sciences. 2022;23. doi: 10.3390/ijms23020986

44. O’Grady GL, Best HA, Sztal TE, Schartner V, Sanjuan-Vazquez M, Donkervoort S, Abath Neto O, Sutton RB, Ilkovski B, Romero NB, et al. Variants in the Oxidoreductase PYROXD1 Cause Early-Onset Myopathy with Internalized Nuclei and Myofibrillar Disorganization. American journal of human genetics. 2016;99:1086–1105. doi: 10.1016/j.ajhg.2016.09.005

45. Caspi T, Straw S, Cheng C, Garnham JO, Scragg JL, Smith J, Koshya AO, Levelt E, Sukumar P, Gierula J, et al. Unique Transcriptome Signature Distinguishes Patients With Heart Failure With Myopathy. Journal of the American Heart Association. 2020;9:e017091. doi: 10.1161/JAHA.120.017091

46. Berndt N, Eckstein J, Wallach I, Nordmeyer S, Kelm M, Kirchner M, Goubergrits L, Schafstedde M, Hennemuth A, Kraus M, et al. CARDIOKIN1: Computational Assessment of Myocardial Metabolic Capability in Healthy Controls and Patients With Valve Diseases. Circulation. 2021;144:1926–1939. doi: 10.1161/CIRCULATIONAHA.121.055646

47. Walker CJ, Schroeder ME, Aguado BA, Anseth KS, Leinwand LA. Matters of the heart: Cellular sex differences. Journal of molecular and cellular cardiology. 2021;160:42–55. doi: 10.1016/j.yjmcc.2021.04.010

48. Lopaschuk GD, Karwi QG, Tian R, Wende AR, Abel ED. Cardiac Energy Metabolism in Heart Failure. Circulation research. 2021;128:1487–1513. doi: 10.1161/CIRCRESAHA.121.318241

49. Sosnowski DK, Jamieson KL, Darwesh AM, Zhang H, Keshavarz-Bahaghighat H, Valencia R, Viveiros A, Edin ML, Zeldin DC, Oudit GY, et al. Changes in the Left Ventricular Eicosanoid Profile in Human Dilated Cardiomyopathy. Front Cardiovasc Med. 2022;9:879209. doi: 10.3389/fcvm.2022.879209

